# *pdb-tools*: a swiss army knife for molecular structures

**DOI:** 10.1101/483305

**Authors:** João P. G. L. M. Rodrigues, João M. C. Teixeira, Mikaël Trellet, Alexandre M. J. J. Bonvin

## Abstract

The pdb-tools are a collection of Python scripts for working with molecular structure data in the PDB format. They allow users to edit, convert, and validate PDB files, from the command-line, in a simple but efficient manner. The pdb-tools are implemented in Python, without any external dependencies, and are freely available under the open-source Apache License at https://github.com/haddocking/pdb-tools/ and on PyPI (https://pypi.org/project/pdb-tools/).

## Introduction

Obtaining and analyzing three-dimensional structures of biological macromolecules, such as proteins or nucleic acids, is often a key step towards understanding their biological function. Of the many file formats used in structural biology to store three-dimensional coordinate data, the Protein Data Bank (PDB) format remains one of the most widely adopted, despite the introduction of a new standard - the mmCIF format - in recent years[1][2]. The PDB format encodes 44 possible different record types in a human-readable flat-text format, with each record having a number of fields with stringent spacing rules in a total of 80 characters. This strictness and complexity of the PDB format make manual editing difficult. As a result, researchers use molecular viewers, e.g. PyMOL[3] and VMD[4], to perform basic editing tasks, such as selecting chains from a structure. More advanced edits, such as changing chain identifiers, deleting specific atoms, or renumbering residues require expertise with scripting languages, whose syntax varies from viewer to viewer. Other edits, such as selecting atomic positions based on occupancy values, are even more challenging to accomplish in molecular viewers, if possible at all. The alternative is for researchers to become proficient in a programming language, such as Python, and use one of its structural bioinformatics libraries, such as BioPython[5] or ProDy[6].

Here, we present pdb-tools, a modular toolkit providing several everyday tasks when handling PDB files, namely downloading, editing, filtering, merging, sorting, and validation, as well as conversion to and from the more recent PDBx/mmCIF file format[2] (Figure 1). In order to shorten the learning curve for new users, all pdb-tools implement a unified command-line interface. Moreover, to support complex editing operations, we designed the toolkit to allow the serial concatenation of several tools in a pipeline without the need to read or write intermediary files. For developers, the coherent architecture simplifies maintenance and development of new tools.

**Figure 1.**
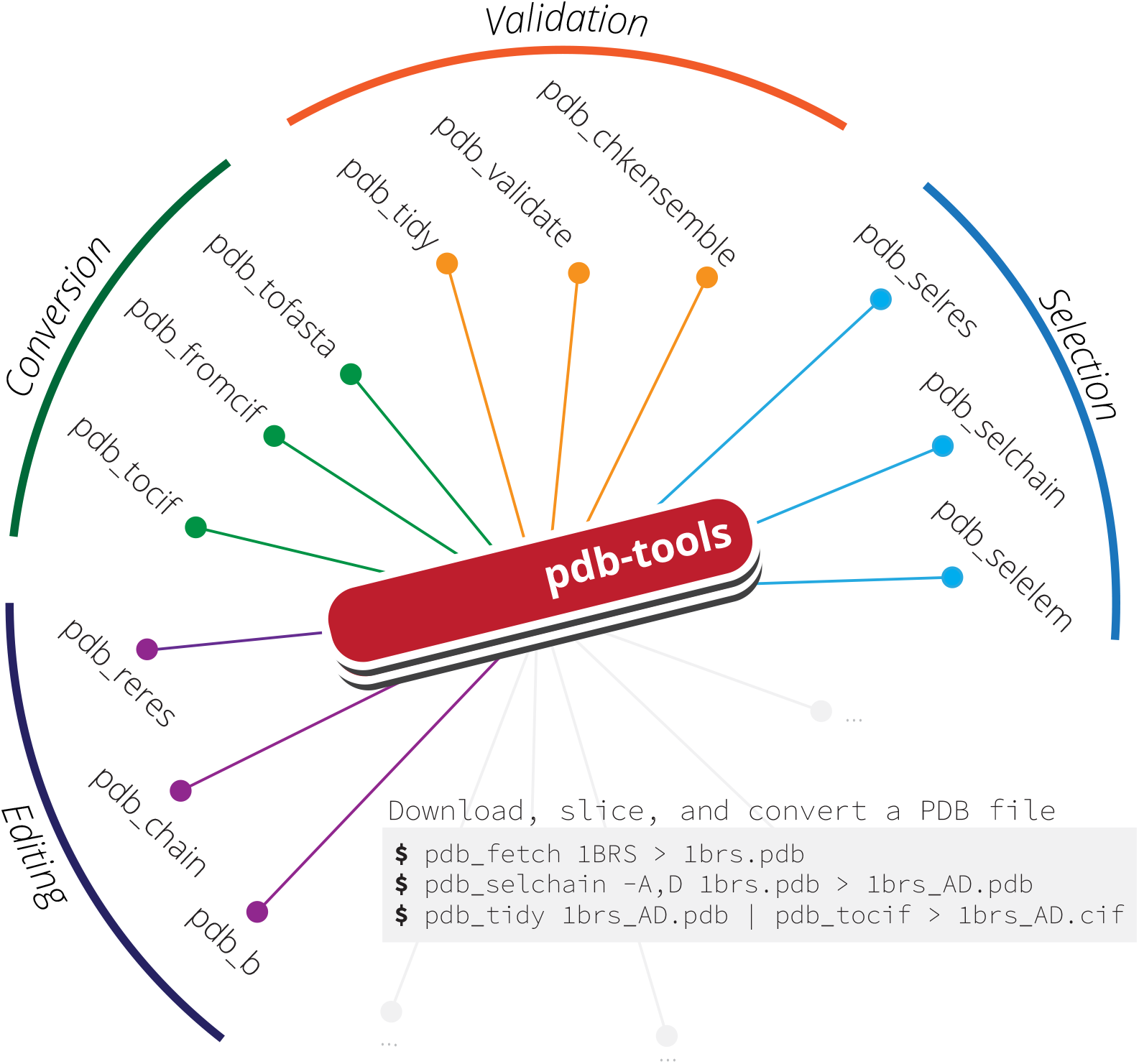
Overview of the tasks supported in pdb-tools and example tools. A common usage example is shown on the gray box insert.

Figure 1. Overview of the tasks supported in pdb-tools and example tools. A common usage example is shown on the gray box insert.

The pdb-tools are written in Python (www.python.org) and are currently developed on GitHub under the open-source Apache License version 2.0. In addition, to simplify the installation procedure for the end-user, the toolkit is also available for download through PyPI (https://pypi.org/project/pdb-tools/). Finally, pdb-tools are also part of the SBGrid initiative[7].

## Methods

### Implementation

The pdb-tools toolkit is implemented in CPython (www.python.org), using only modules of the standard library, which allows us to support a wide range of Python versions (2.7 and all of the 3.x series). The tools make use of Python generator expressions to improve performance and memory usage. This feature, along with support for streaming input and output data, allows users to create complex pipelines by chaining together different tools (e.g. format conversion followed by chain selection followed by editing followed by validation).

As a result, using ‘pdb_reres’ to renumber the residues of a structure with 64,606 atoms takes ~0.17 seconds (on an Intel i7-7500U CPU) and ~10 MB of memory. Merging eight copies of the same structure using ‘pdb_merge’ took ~0.74 seconds and ~18 MB of memory. These performance indicators make pdb-tools particularly useful when combined with shell scripting for batch processing entire collections of PDB files.

To aid the maintenance and development of pdb-tools, we wrote approximately 250 separate unit tests that provide 81% coverage for the tools and their possible usage options. In addition, each commit is checked for coding style against the PEP8 style guide, using flake8. Besides hosting the source code on GitHub, we use setuptools to package and distribute pdb-tools through PyPI. Further, we use semantic versioning to indicate compatibility between releases with the same major version. The code is tested continuously using Travis CI and AppVeyor webhooks on GitHub, which monitor incoming pull requests and any commits to the master branch. We test on virtual instances of Windows 10 and Linux 16.04 LTS, under Python 2.7, 3.6, and 3.7. Nevertheless, the pdb-tools should run on most UNIX derivatives and Windows versions, provided a supported version of Python in installed.

Concerning documentation, each tool contains a self-contained help string, describing its purpose, command-line interface, and usage examples. This help text is available by running the tool without any argument. Additional documentation on the usage and installation of pdb-tools is available online on the project’s webpage (www.bonvinlab.org/pdb-tools) and in the README file accompanying the source code.

Finally, we develop pdb-tools as a community open-source project on GitHub and encourage contributions of any kind. To date, the project has more than 30 forks and has been starred by over a dozen users on GitHub. To help and promote these contributions, and based on our experience with building pdb-tools collaboratively, we have documents setting coding best-practices, contribution guidelines, and a code of conduct for developers on our GitHub repository. In addition, we make use of and encourage users to contribute to our issue tracker, which documents our discussions and development decisions publicly.

## Use Cases

The pdb-tools follow a ‘one tool, one job’ design philosophy, where each tool performs a simple operation on an input file, and more complex operations can be achieved by chaining different tools together. Over the years, the pdb-tools have been used in both research and educational contexts, some of which we highlight below.

### Making selections in PDB files

Molecular viewers, such as PyMOL or VMD, are the tools of choice for making selections of specific chains or residue ranges in PDB files. When handling more than one structure, programming libraries such as Biopython or Prody are better suited for the task. The selection tools includes in pdb-tools offer a simple solution to make complex selections.

~~~
$ pdb_fetch 1ctf | pdb_keepcoord | pdb_reres > 1ctf.pdb
$ pdb_selres -1:30 1ctf.pdb | pdb_selatom -CA > 1ctf_CA.pdb
$ head 1ctf_CA.pdb
ATOM     2  CA  GLU A  1    17.706  17.982 -14.905 1.00 16.74   C
ATOM     11 CA  PHE A  2    17.509  14.262 -14.184 1.00 13.24   C
... [truncated]
$
~~~

Importantly, users can use shell scripting to efficiently process large collections of structures. Those using UNIX-based systems can also make use of the parallel utility to distribute processing over several cores.

~~~
$ for pdbf in $( ls *.pdb )
$ do
$ pdb_selchain -A $pdbf | pdb_delelement -H | pdb_tidy > ${pdbf}_A_noH.pdb
$ done
~~~

### Preparing input files for molecular modelling using HADDOCK

HADDOCK is an integrative modeling software for protein interactions[8], whose web server provides a user-friendly interface that oversaw over 200.000 job submissions to date. The simplest of submissions requires two structures, in PDB format, that cannot have more than one chain, cannot have overlapping residue numbers, and cannot have alternate locations for any atom. Often, users submitting jobs to the HADDOCK web server are faced with error messages because of such formatting requirements. Using pdb-tools, producing a HADDOCK-compliant PDB file is straightforward, as we demonstrate below.

~~~
$ pdb_fetch 1brs | pdb_keepcoord > 1brs.pdb # 6 chains
$ pdb_selchain -A,D 1brs.pdb | pdb_delhetatm | pdb_selaltloc > 1brs_AD.pdb
$ pdb_selchain -A 1brs_AD.pdb | pdb_tidy > 1brs_A.pdb
$ pdb_selchain -D 1brs_AD.pdb | pdb_tidy > 1brs_D.pdb
~~~

### Extracting the aminoacid/nucleotide sequence of a PDB file

Although PDB files may include SEQRES records that store the sequence of the construct used for structure determination, the final structure does not necessarily include all the amino acids or nucleotides. This discrepancy is particularly important when building alignments for homology modeling. To this end, we include in pdb-tools a PDB to FASTA converter that extracts the sequence of the structure directly from the ATOM/HETATM records. The user can specify if the entire structure should be exported as a single FASTA sequence record, or divided by chain, using the -**multi** option.

~~~
$ pdb_fetch 1brs | pdb_tofasta
>PDB|ABCDEF
INTFDGVADYLQTYHKLPDNYITKSEAQALGWVASKGNLADVAPGKSIGGDIFSNREGKL
PGKSGRTWREADINYTSGFRNSDRILYSSDWLIYKTTDHYQTFTKIRAQVINTFDGVADY
... [truncated]
$ pdb_fetch 1brs | pdb_tofasta -multi
>PDB|A
VINTFDGVADYLQTYHKLPDNYITKSEAQALGWVASKGNLADVAPGKSIGGDIFSNREGK
LPGKSGRTWREADINYTSGFRNSDRILYSSDWLIYKTTDHYQTFTKIR
>PDB|B
AQVINTFDGVADYLQTYHKLPDNYITKSEAQALGWVASKGNLADVAPGKSIGGDIFSNRE
GKLPGKSGRTWREADINYTSGFRNSDRILYSSDWLIYKTTDHYQTFTKIR
... [truncated]
~~~

### Converting to and from mmCIF format

Due to several limitations with the PDB file format, the Protein Data Bank recently introduced the new mmCIF format. To support this decision, we include in pdb-tools converters to and from the mmCIF format:

~~~
$ pdb_fromcif 1brs.cif | pdb_selchain -A | pdb_tocif > 1brs_A.cif
~~~

## Summary

The pdb-tools provide a command-line interface to edit three-dimensional molecular structures in the PDB format. Since it is written in Python, without any third-party dependencies, the toolkit is available on Linux, Mac OS, and Windows and supported on both Python 2 and 3. The source code is hosted on GitHub and licensed under the open-source Apache License version 2.0, to encourage contributions from the community, and available for installation through PyPI and SBGrid. As such, the pdb-tools are an efficient, portable, and extremely flexible toolkit for both starting and experienced researchers in structural biology.

## Software availability

1. https://haddocking.github.io/pdb-tools/
2. https://pypi.org/project/pdb-tools/
3. https://sbgrid.org/software/titles/pdb-tools
4. https://github.com/haddocking/pdb-tools
5. https://github.com/haddocking/pdb-tools/releases/tag/2.0.0-beta.5
6. https://github.com/haddocking/pdb-tools/releases/tag/2.0.0-beta.5
7. Apache License, version 2.0

## Author contributions

All authors contributed and validated code and participated in the drafting and revisions of the manuscript. J.P.G.L.M.R and A.M.J.J.B supervised the research.

## Acknowledgments

The authors acknowledge the members of the Bonvin group for fruitful discussions and suggestions of new tools over the years.

